# Do fungi look like macroparasites? Quantifying the patterns and mechanisms of aggregation for host-fungal parasite relationships

**DOI:** 10.1101/2024.08.29.609018

**Authors:** Sarah A.R. Schrock, Jason C. Walsman, Joseph DeMarchi, Emily H. LeSage, Michel E.B. Ohmer, Louise A. Rollins-Smith, Cheryl J. Briggs, Corinne L. Richards-Zawacki, Douglas C. Woodhams, Roland A. Knapp, Thomas C. Smith, Célio F.B. Haddad, C. Guilherme Becker, Pieter T.J. Johnson, Mark Q. Wilber

**Affiliations:** School of Natural Resources, University of Tennessee Institute of Agriculture, Knoxville, TN, USA; Ecology, Evolution, and Marine Biology, University of California, Santa Barbara, CA, USA; Biology Department, Skidmore College, Saratoga Springs, NY, USA; Department of Biology, University of Mississippi, University, MS, USA; Department of Pathology, Microbiology, and Immunology, Vanderbilt University School of Medicine, Nashville, TN, USA; Department of Biological Sciences, Vanderbilt University, Nashville, TN, USA; Department of Biological Sciences and Pymatuning Laboratory of Ecology, University of Pittsburgh, PA, USA; Department of Biology, University of Massachusetts, Boston, MA, USA; Sierra Nevada Aquatic Research Laboratory, University of California, Mammoth Lakes, CA, USA; Earth Research Institute, University of California, Santa Barbara, CA, USA; Department of Biodiversity and Aquaculture Center (CAUNESP), Universidade Estadual Paulista, Rio Claro, SP, Brazil; Department of Biology, The Pennsylvania State University, University Park, PA, USA; One Health Microbiome Center, Center for Infectious Disease Dynamics, Ecology Institute, Huck Institutes of the Life Sciences, The Pennsylvania State University, University Park, PA, USA; Ecology and Evolutionary Biology, University of Colorado Boulder, Boulder, CO, USA

**Author notes:** **Corresponding author**: Sarah A.R. Schrock, School of Natural Resources, University of Tennessee Institute of Agriculture, Knoxville, TN, USA.

## Abstract

Most hosts contain few parasites, whereas few hosts contain many. This pattern, known as aggregation, is well-documented in macroparasites where parasite intensity distribution among hosts affects host-parasite dynamics. Infection intensity also drives fungal disease dynamics, but we lack a basic understanding of host-fungal aggregation patterns, how they compare to macroparasites, and if they reflect biological processes. To address these gaps, we characterized aggregation of the fungal pathogen *Batrachochytrium dendrobatidis* (Bd) in amphibian hosts. Utilizing the slope of Taylor’s Power Law, we found Bd intensity distributions were more aggregated than macroparasites, conforming closely to lognormal distributions. We observed that Bd aggregation patterns are strongly correlated with known biological processes operating in amphibian populations, such as epizoological phase—invasion, post-invasion, and enzootic—and intensity-dependent disease mortality. Using intensity-dependent mathematical models, we found evidence of evolution of host resistance based on aggregation shifts in systems persisting with Bd following disease-induced declines. Our results show that Bd aggregation is highly conserved across disparate systems and is distinct from aggregation patterns in macroparasites, and contains signatures of potential biological processes of amphibian-Bd systems. Our work lays a foundation to unite host-fungal dynamics under a common theoretical framework and inform future modeling approaches that may elucidate host-fungus interactions.

## Introduction

One of the few general laws of parasitology is that many hosts have few parasites, and few hosts have many parasites [1]. Known as ”aggregation”, this pattern has important implications for the dynamics of host-parasite systems and our ability to infer the dominant processes operating within them [2; 3; 4]. For example, some macroparasites can cause intensity-dependent parasite-induced mortality, and the severity of this process can be reflected in the intensity distribution of parasites across hosts [5; 6]. In wildlife-macroparasite systems, such as nematodes, trematodes, and ectoparasitic arthropods, the nature of aggregation has been extensively quantified [7; 8]: the distribution of macroparasites among hosts is often well-described by a negative binomial distribution, and variance-to-mean relationships are significantly different from Poisson expectations. While we have long been able to quantify the intensity of macroparasites (e.g., by counting parasites following dissection), we can now also quantify infection intensity of microparasites through the broad application of modern molecular techniques. Microparasites are organisms such as bacteria, viruses, protozoa, and fungi that have high replication rates within a host and often induce host immune responses [9]. While studies on microparasites now regularly report quantitative measures of infection (e.g., viral titers or fungal intensity within a host), we have few baseline expectations regarding what the intensity distributions of microparasites look like and the mechanisms shaping them.

Here, we focus on fungal parasites. Fungal parasites are a global threat to wildlife populations [10]: e.g., *Batrachochytrium dendrobatidis* (Bd), *B. salamandrivorans*, *Ophidiomyces ophiodiicola*, and *Pseudogymnoascus destructans* have led to dramatic declines and extinctions in hundreds of wildlife species [11; 12; 13; 14]. Like macroparasites, animals infected with fungal parasites suffer intensity-dependent parasite-induced mortality [15; 16]. This means that accounting for the distribution of fungal parasite intensity within a population is critical for predicting population-level outcomes following fungal invasion [17; 18]. However, despite modeling work increasingly accounting for fungal infection intensity [17; 19], we still lack a general understanding of the quantitative patterns of aggregation in host-fungal parasite systems. Quantifying these patterns is important because i) different levels of aggregation change system dynamics and can significantly affect model predictions [20; 21] and ii) patterns in fungal intensity distributions may reflect dominant mechanistic processes structuring the host-parasite system [22]. The latter is particularly important for parasites like Bd where cryptic disease-induced mortality may drive ongoing declines [23], but detecting these declines is difficult. Aggregation patterns in fungal intensity distributions could potentially provide a mechanism to detect signatures of disease-induced mortality, as has been done in host-macroparasite systems [6].

Describing the distribution of fungal parasite intensity requires a different statistical and conceptual treatment than traditional macroparasite models. Macroparasite infection intensity is typically described by parasite counts—in other words, how many parasites are found within a host, ranging from zero to some large number. As such, macroparasite counts are discrete and can be described by distributions such a Poisson or negative binomial distribution [8]. In contrast, fungal parasite intensity is typically quantified by molecular approaches such as quantitative PCR [qPCR; 24]. The qPCR technique measures the amount of a specific DNA sequence in a sample by amplifying the sequence while simultaneously detecting and quantifying the fluorescence of the product in real-time as the reaction proceeds. Because the amount of fluorescence generated is directly proportional to the amount of starting DNA, qPCR values correlate with fungal intensity. The resulting measurement of ”infection intensity” is a continuous variable ranging from zero to some arbitrarily large number.

Using infection intensity as a continuous quantity computed by qPCR presents two methodological challenges. First, qPCR measures of infection intensity are subject to substantial measurement error [25; 26]. Measurement error can come when the sample of infection intensity is collected (e.g., the skin of amphibians infected with Bd are swabbed) or when the sample is processed with qPCR [26]. For example, the qPCR process often fails to detect very low quantities of genetic material and can miss low levels of infection [27; 26]. Generally, increasing the noise in a sample due to measurement error might decrease our ability to detect biological signals. Thus, we might expect measurement error to play a more significant role in affecting the patterns of aggregation in fungal intensity distributions than typical macroparasite distributions, obscuring mechanistic signatures of host-parasite processes on fungal intensity distributions.

Second, discrete distributions that are typically used to describe macroparasite counts are not technically applicable to continuous molecular infection intensity data. In amphibian-Bd systems, there has been some previous discussion on reasonable assumptions for the distribution of infection intensity [particularly with regards to the random component of generalized linear models; 25] and how approximating a continuous random variable with a discrete random variable [e.g., using a negative binomial distribution to describe infection intensity 25] can affect the conclusions one draws. However, there has been no systematic examination of the distribution that most consistently describes observed amphibian-Bd distributions or parasitic fungal distributions more broadly. As we continue to develop models for predicting the dynamics of fungal outbreaks, a systematic quantification of the nature of fungal intensity distributions can help direct these modeling efforts, as it has done in traditional macroparasite systems [7; 8].

In addition to these statistical differences, there are key biological differences between fungal parasites and macroparasites that may affect observed patterns of aggregation. Fungal parasites grow within/on a host leading to increases in infection intensity. Typically (though not always), macroparasite infections increase in intensity through ”immigration processes” rather than ”birth processes”—hosts repeatedly encounter parasites in the environment which leads to an accumulation of parasites. Birth processes such as the within host reproduction of parasites are known to increase the aggregation of macroparasite distributions [28; 29]. An initial expectation might be that fungal distributions are typically more aggregated than macroparasite distributions. However, this prediction is complicated by the speed and mode of transmission of fungal parasites, which can be faster than many macroparasites. For example, Bd can complete its life cycle in four to ten days, whereas a trematode parasite with multiple intermediate hosts might take months to complete its life cycle [30; 31]. This could lead to faster spread, more homogenization, and lower levels of aggregation for fungal parasites like Bd compared to macroparasites.

Here, we utilized 56,912 skin swab samples from 93 amphibian species to ask two main questions: (1) What is the general structure of these fungal intensity distributions, and (2) do they reflect biological processes? First, we examined whether we see aggregation in host-Bd systems, how these patterns compare to those of macroparasites, and what statistical distribution best describes these fungal intensity distributions. We hypothesized that i) fungal distributions will be aggregated, ii) they will show higher levels of aggregation than most macroparasite distributions, and iii) they will generally conform to a lognormal distribution. Our prediction of a lognormal distribution stems from theoretical work showing that lognormal distributions robustly describe population densities subject to demographic and environmental stochastic-ity, as well as measurement error [32]. To address our second question, we compared aggregation patterns among amphibian-Bd systems in different epizoological states (e.g., invasion, post-invasion, and enzootic) to see if they reflect underlying biological processes. To complement data analysis, we employed an integral projection model to gain insight into the possible mechanisms driving the observed aggregation patterns. Given intensity-dependent disease dynamics in amphibian-Bd systems, we expected reduced aggregation in populations experiencing significant disease-induced mortality, such as those in post-invasion, epizootic states. Similarly, we expected disease-induced mortality to be a critical model parameter in reproducing these patterns.

## Materials and Methods

### Amphibian-Bd infection intensity data

We analyzed four datasets of Bd infection intensities (henceforth ”intensity” or ”load”) obtained from amphibian skin swabs collected in the field. Bd loads were obtained through DNA extraction and qPCR, which detects the number of genomic equivalents or ITS1 copy number of Bd on amphibian skin. These procedures were standardized within but not across datasets. As such, it is important to note that our analysis does not aim to compare absolute values of fungal intensities across datasets or even among disparate sites within datasets. Variations in techniques between labs and calculations of Bd intensity (e.g. multiplying by different scaling factors) as well as differences in ITS copy numbers for different strains of Bd in different sites [e.g., 33] could make comparisons challenging. Instead, we use measures of aggregation (described below) that are scale invariant, thus providing robust measures to analyze aggregation patterns. However, if individuals of the same species of amphibian in the same site in the same season are co-infected with different strains of Bd that vary in their ITS1 copy number [e.g., 34], then the aggregation metrics we estimate could suffer bias.

The first dataset we included was from Brazil (henceforth the Brazil dataset) which contained 4,365 swabs from 41 amphibian species collected primarily within the state of São Paulo (see Fig. S1 for general locations of research sites for all datasets). Our second dataset comes from the East Bay region of California (henceforth the East Bay dataset) and contains 10,490 swabs from 11 host species. The third dataset contains 12,457 Bd swabs from amphibians collected from 2016-2019 on 43 amphibian species across 31 research sites in four states—Louisiana, Pennsylvania, Tennessee, and Vermont. Although collected across a wide geographical range, swabs from this study were all processed at a centralized location using a consistent methodology. Therefore, we will refer to this dataset broadly as the Eastern US dataset. Our final dataset is from the Sierra Nevada mountains of California (henceforth the Sierra dataset) and contains 29,600 samples collected from mountain yellow-legged frogs (MYL frogs; composed of sister species *Rana muscosa* and *Rana sierrae*) at high elevation lakes, ponds, and wetlands.

Samples within each dataset were grouped based on host species, life stage (larva, subadult, or adult), research site, season (Brazil: Wet or Dry; East Bay and Sierra: Summer; Eastern US: winter, spring, summer, or fall), and year (see Table S1 for more detailed composition of each dataset). Moving forward, we will refer to a particular combination of species, life stage, research site, season, and year as a ”group”. Examining specific ”groups” allows us to quantify the patterns of Bd aggregation in a biologically relevant temporal period at a particular location. In total, the Brazil dataset had 109 candidate groups for analysis, East Bay had 714, Eastern US had 391, and the Sierra had 647.

### Question 1a: Are fungal intensity distributions aggregated and how do these compare with aggregation patterns in macroparasite systems?

To address this question, we analyzed aggregation in the fungal intensity distributions using Taylor’s Power Law (TPL) which relates the log mean and log variance in fungal intensity, calculated for each group. This metric allows for direct comparison to the macroparasite literature. Specifically, we focused on the slope of TPL as a metric of aggregation, where a greater slope indicates greater aggregation [28; 3]. Across all datasets, we only included groups with at least three infected individuals, yielding 961 groups across all four datasets (Table S1).

We first fit a linear regression to the log mean vs. log variance relationship for each of the four datasets and calculated the slope. We compared the slopes to the empirical relationship previously seen in macroparasite populations (slope=1.55, 95% Confidence Interval: [1.48,1.62]) [7], as well as a Poisson distribution (mean-variance slope equal to 1), which is generally considered the null distribution in many host-macroparasite studies [2]. However, the continuous nature of Bd load data also suggests considering the alternative null with a TPL slope of 2. A baseline of TPL slope of 2 has been used to describe the aggregation of free-living organisms in space and time [35; 36]. Moreover, given our expectation of a lognormal distribution of Bd intensity across hosts, we would expect a TPL slope of 2 based on the simple definitions of the mean and variance for a lognormal distribution. Note that for this analysis, the log mean and the log variance for each group was computed using both infected and uninfected individuals, consistent with macroparasite studies. Second, to explore variability in the slope of TPL across the 36 species with sufficient sampling, we ran a linear mixed effect model (i.e., Gaussian error) with random effects of amphibian species and sub-region on the intercept and slope. Specifically, the model we fit was log(variance) *∼* log(mean) + (1 + log(mean)*|*subregion) + (1 + log(mean)*|*species), where subregion was a factor with the following levels: East Bay, Sierra, Pennsylvania, Tennessee, Vermont, Louisiana, and Brazil. We then examined the species-specific TPL slopes and compared them to the macroparasite slope from Shaw and Dobson (1995) [7].

### Question 1b: What distribution best describes fungal intensity distributions?

To characterize the shape of fungal intensity distributions conditional on infection, we considered continuous distributions of nonnegative real numbers: gamma, exponential, lognormal, and Weibull. We did not consider Poisson and negative binomial distributions because fungal intensity, as assessed using qPCR, is a continuous measure. Although qPCR results can be transformed into integer values and analyzed using standard generalized linear models [37], we opted to keep the data on the continuous scale, consistent with previous models [19]. Each of the continuous distributions can capture a strong right skew in intensity distributions, consistent with canonical patterns in host-macroparasite systems. The gamma distribution is the continuous analog to the negative binomial distribution, a distribution that describes many macroparasite populations [8]. Similarly, the exponential is a special case of the gamma distribution that is represented by only one parameter and is analogous to the discrete geometric distribution which has been proposed as a potential null distribution in host-macroparasite systems [38; 39]. Lognormal distributions are found throughout natural systems empirically and theoretically [32] and are representative of nonnegative metrics with relatively low means but large variance. Finally, we considered the Weibull distribution which is typically used to model ”time-to-failure” or survival analyses but has been used to describe macroparasite aggregation data [40].

For this analysis, we only considered groups with at least 10 infected individuals to ensure we had power to distinguish between competing distributions. This resulted in 525 groups. We used the fitdistrplus package in R to fit exponential, lognormal, Weibull, and gamma distributions using maximum likelihood estimation (MLE) or moment matching estimation, if the MLE model would not converge. We compared Akaike information criterion (AIC) values across distributions to find the best predictive model, assuming no notable difference in performance when AIC values were within +/- 2.

### Question 2: Do patterns of aggregation in Bd intensity reflect biological processes, such that there are quantifiable differences in aggregation between epizoological states?

To address this question, we used a metric that can be applied to a single group (unlike TPL) known as Poulin’s Discrepancy Index, or simply Poulin’s D [41; 4]. Poulin’s D is bounded from 0 to 1 and is a proportional measure of the difference between an observed distribution and a uniform distribution. A higher value indicates greater discrepancy from a uniform distribution and is suggestive of higher aggregation. The equation for Poulin’s D is 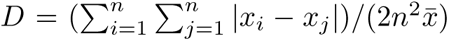, where *x* is the fungal load of host *i* or *j*, *n* is the total number of hosts, and 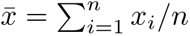 [we use the equation given in 42, which is the Gini index]. We also calculated the coefficient of variation (CV) on the natural scale and other related metrics—log_10_-transformed CV on the natural scale and CV on the log_10_ scale—which should provide comparable results to Poulin’s D [42]. We calculated CV on the log_10_-transformed data to determine if trends remained similar on different scales. When calculating our aggregation metrics, we excluded uninfected individuals to remove the effect of prevalence on the observed patterns. We only included groups that had at least two infected individuals—the minimum number for a meaningful value of our metrics. We also explored only including groups with a minimum of 10 infected individuals, and our results were unchanged.

For this question, we focused on the Sierra dataset. Of the datasets used in this study, the Sierra dataset is unique because for many southern populations in the Sierra Nevada, we know when Bd invaded, when epizootics ensued, and when populations declined [43; 15; 44]. Moreover, for more northern populations, such as those in Yosemite National Park, we know that populations are past the invasion-epizootic-declining phase and are persisting enzootically with Bd [45; 17]. Thus, we have three clearly definable epizoological phases for MYL-Bd populations in the Sierra Nevada: 1) invasion stage [when Bd prevalence is less than 50% in a population; 46] 2) post-invasion phase (consisting of epizootic host declines or recent declines) and 3) enzootic phase (Bd invaded before the early 2000’s and amphibian populations are persisting in the presence of Bd). Moreover, from targeted field surveys and laboratory experiments, we know that there is strong intensity-dependent mortality in MYL frogs [15; 19]. If patterns of Bd aggregation contain information about intensity-dependent mortality, we would expect a notable reduction in Bd aggregation for higher mean infection intensity in MYL frog populations [3]. In other words, mortality in highly infected individuals would effectively reduce the tail of the right-skewed distribution characteristic of aggregated populations, thereby decreasing aggregation.

To explore signatures of epizoological phase on Bd aggregation, we first plotted each metric—Poulin’s D, CV, log CV, and CV of log-scale data—against mean log_10_-transformed Bd intensity and asked whether populations in known epizoological phases clustered in mean intensity-aggregation space (henceforth intensity-aggregation space) and whether there were notable reductions in aggregation at high infection intensities (note that epizoological phases were determined independently of aggregation or mean infection intensity). We used beta regression [4] to test for a quadratic effect of mean infection intensity on aggregation metrics, where a strong quadratic effect is indicative of aggregation being reduced at high infection intensity.

Finally, to better understand how mechanisms such as intensity-dependent mortality and epizoological phase could theoretically affect patterns of aggregation in host-fungal systems, we adapted an Integral

Projection Model (IPM) that has been previously developed for amphibian-Bd systems [19]. In short, IPMs provide an approach for modeling intensity-dependent infection dynamics of host-fungal interactions by specifically modeling the entire distribution of fungal intensities within a population (see supplementary material for more detail). Hosts are born uninfected, and in the absence of disease, the host population grows logistically toward a carrying capacity. In one time step of the model, hosts may become infected by encounter with environmental pathogens and gain some initial log number of parasites (infection load). Parasites grow within hosts, with some stochasticity, toward a within-host carrying capacity. In each time step, infected hosts have a probability of recovery from infection and a probability of survival, both of which decline with infection load. Infected hosts shed parasites back into the environment proportional to the number of parasites they currently hold. We simulated disease invasion for one year to represent the effects of disease spread without host evolution. We then added simulations where we included multiple host genotypes with different traits to simulate evolution over 30 years (a relevant timescale for the MYL-Bd system). Specifically, we focused on host evolution of resistance that lowers pathogen growth rate, an important mechanism in the MYL-Bd system [45; 37]. We performed simulations at parameter values from laboratory experiments for the MYL-Bd system (see Table S2 in supplementary material) and then explored how varying certain parameters impacted the intensity-aggregation patterns in our simulations. We calculated the same four aggregation metrics in our simulations as were calculated from field data to determine intensity-aggregation patterns. We did this to investigate the patterns that could emerge in intensity-aggregation space for the different metrics and if they are indicative of specific biological mechanisms.

## Results

### Question 1a: Are fungal intensity distributions aggregated and how do these compare with aggregation patterns in macroparasite systems?

Based on TPL, Bd showed a greater degree of aggregation compared to macroparasites (Fig. 1). The slopes of TPL across the groups for each dataset ranged between 1.90, 95% CI [1.86-1.94] (Sierra) and 2.06, 95% CI [1.99-2.12] (Brazil), which are all significantly higher than the macroparasite slope given by Shaw and Dobson (1.55, 95% CI [1.48-1.62])[7] (Fig. 1). Therefore, the variance of Bd infection intensity increases to a greater degree with respect to average fungal load than many macroparasites.

**Figure 1:**
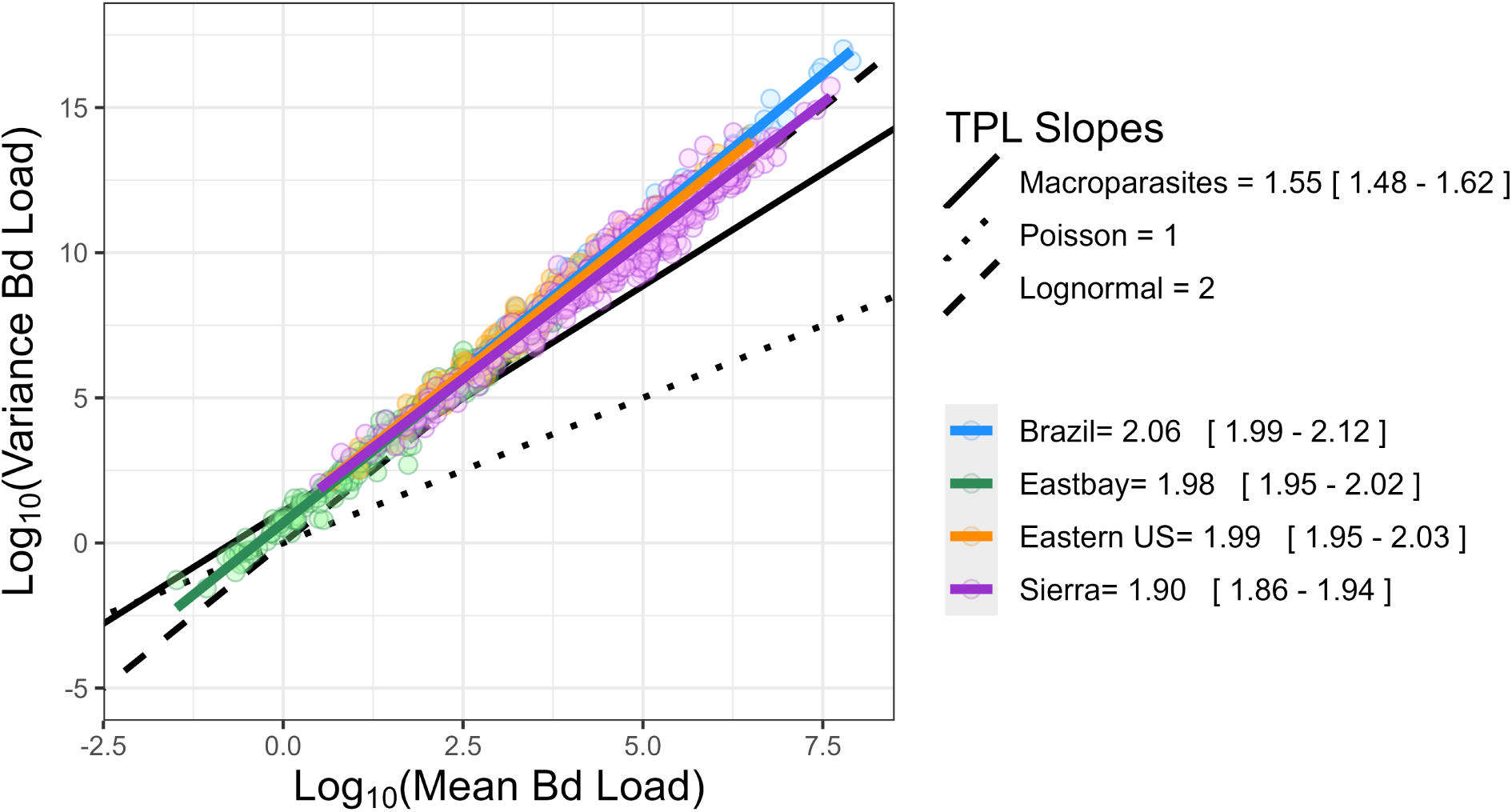
The relationship of log mean and log variance of fungal intensity for all groups. Regression lines were fit to each dataset. Slopes and their 95% confidence intervals are provided in the legend. The solid black line represents the slope that is typically seen in macroparasites (1.55, 95% CI [1.48-1.62])[7]. The dotted line with a slope of 1 is expected in a Poisson distribution (null distribution for macroparasites) and the dashed line with a slope of 2 is expected for a lognormal distribution.

We examined how the slopes of TPL varied among amphibian species and life stages. While we found significant variation in the slope of TPL among species (including slope as a random effect among species yielded a better predictive model than a model without a species-level random effect of slope: ΔAIC=13.6), Bd was more aggregated on all amphibian species than for many macroparasites (see Fig. S2). We found a similar pattern across host life stage, showing slopes greater than that of macroparasites. However, the slopes for each life stage were statistically distinct: larval (1.85, 95% CI [1.82-1.89]), subadult (1.93, 95% CI [1.91-1.95]), and adult (2.01, 95% CI [1.98-2.04]).

### Question 1b: What distribution best describes fungal intensity distributions?

Of the distributions that we fit to the Bd-positive data, the lognormal model consistently performed better than the others, as determined by comparing AIC scores (Fig. 2). Assuming models perform equally well if AIC scores are within 2 units of each other, over half of the groups (57.3%) were well-described by multiple distributions. The lognormal performed best or just as well as another model in 76.7% of the groups, the Weibull in 58.8%, the gamma in 35.2%, and the exponential in 25.7%. The lognormal model also fit 38.0% of groups better (*>* 2 AIC units) than any of the other models. Whereas, the Weibull, gamma, and exponential models performed better than all others in 4.4%, 0.4%, and 0% of the groups, respectively.

**Figure 2:**
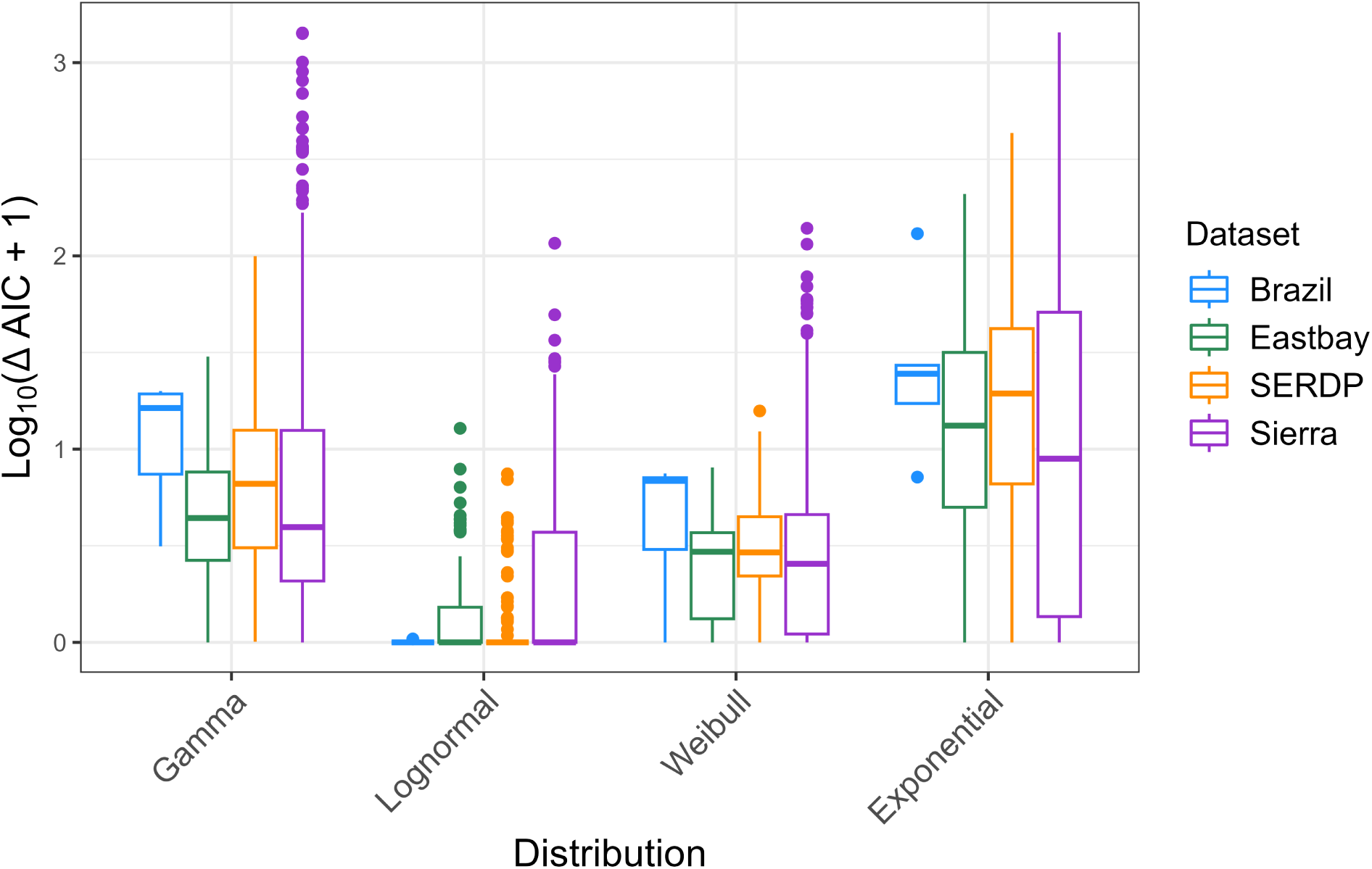
Comparison of log_10_(Δ*AIC* + 1) values across the four continuous distributions that were fit to the fungal intensity data for the 525 amphibian groups across four datasets. Each data point was a group of amphibians that had at least 10 infected individuals.

With the lognormal model outperforming the other distributions, we sought to determine if the lognormal is objectively a good fit to the data. We used a Shapiro-Wilk’s test of normality on the log-transformed data, after adjusting the p-values for multiple tests to account for false discovery rate (using the p.adjust function in R with method fdr). For 96.7% of sampled groups, we fail to reject the null hypothesis that the data follows a normal distribution (Fig. S3, at an adjusted significance level of *α* = 0.05). Cognizant that failure to reject the null is not proof of the null, we conclude there is not strong evidence that distributions deviate from a lognormal distribution.

### Question 2: Do patterns of aggregation in Bd intensity reflect biological processes, such that there are quantifiable differences in aggregation between epizoological states?

#### Empirical results

To gain mechanistic intuition on the broader results in this section, we first examined seven specific populations from the Sierra dataset that 1) were repeatedly surveyed during Bd invasion and declines and 2) had sufficient samples of infected adults or subadults at a minimum of three time points to compute Poulin’s D (*n ≥* 2). Fig. 3A shows the abundance trajectory of adult frogs in these populations through time, including the well-known pattern of dramatic population declines following Bd invasion. In Fig. 3B, we plot these same populations in intensity-aggregation space and see a consistent counterclockwise pattern emerge. Upon invasion, mean infection intensity is low, and aggregation is low. Once the population transitions to the post-invasion phase, mean intensity is high, but aggregation remains relatively low. As the population progresses through the epizootic, mean intensity declines and aggregation increases. These patterns suggest that there is a signature of epizoological phase on observed patterns of aggregation.

**Figure 3:**
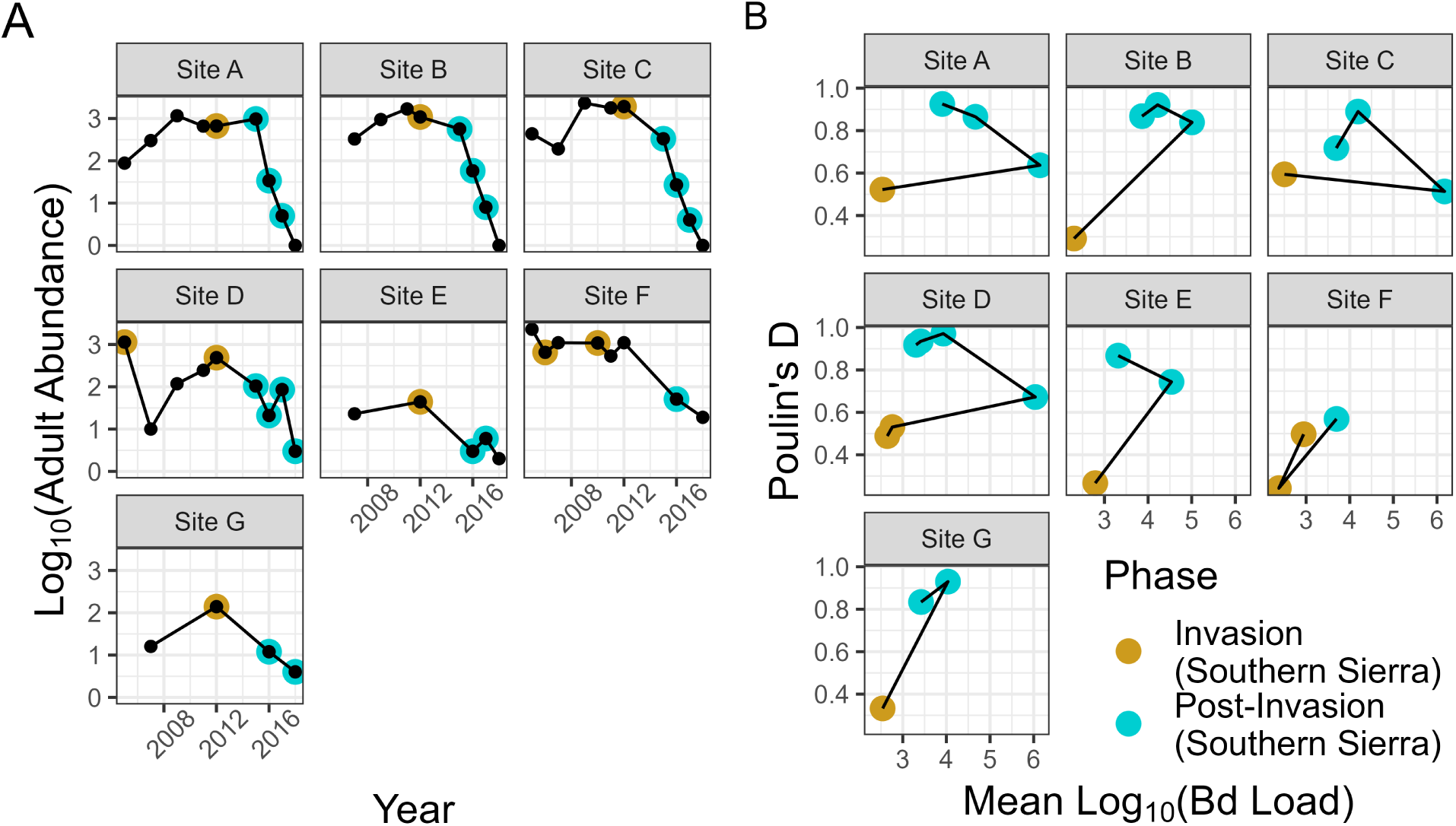
**A.** The adult abundance trajectories of seven focal MYL frog populations through time. Black points show each time the population was surveyed for abundance and Bd and colored points indicate when a sufficient number of infected individuals (*n ≥* 2) were sampled to compute Poulin’s D, with a higher value indicating more aggregation. The invasion phase was delineated when prevalence was less than 0.5, following Wilber et al. (2022) [46]. **B.** The same seven populations with trajectories plotted in intensity-aggregation space. The colored dots in B. correspond to the same colored dots in A.

To examine this pattern more broadly, we plotted 313 Sierra groups in intensity-aggregation space and observed a strong clustering of invasion, post-invasion, and enzootic groups (Fig. 4A-D) that was consistent with what we saw in our seven focal populations with time series data (Fig. 3B). Namely, the invasion stage was characterized by low mean intensity and low aggregation, the post-invasion phase was characterized by medium to high intensity and high to low aggregation, and the enzootic phase was characterized by intermediate mean intensity and high aggregation.

**Figure 4:**
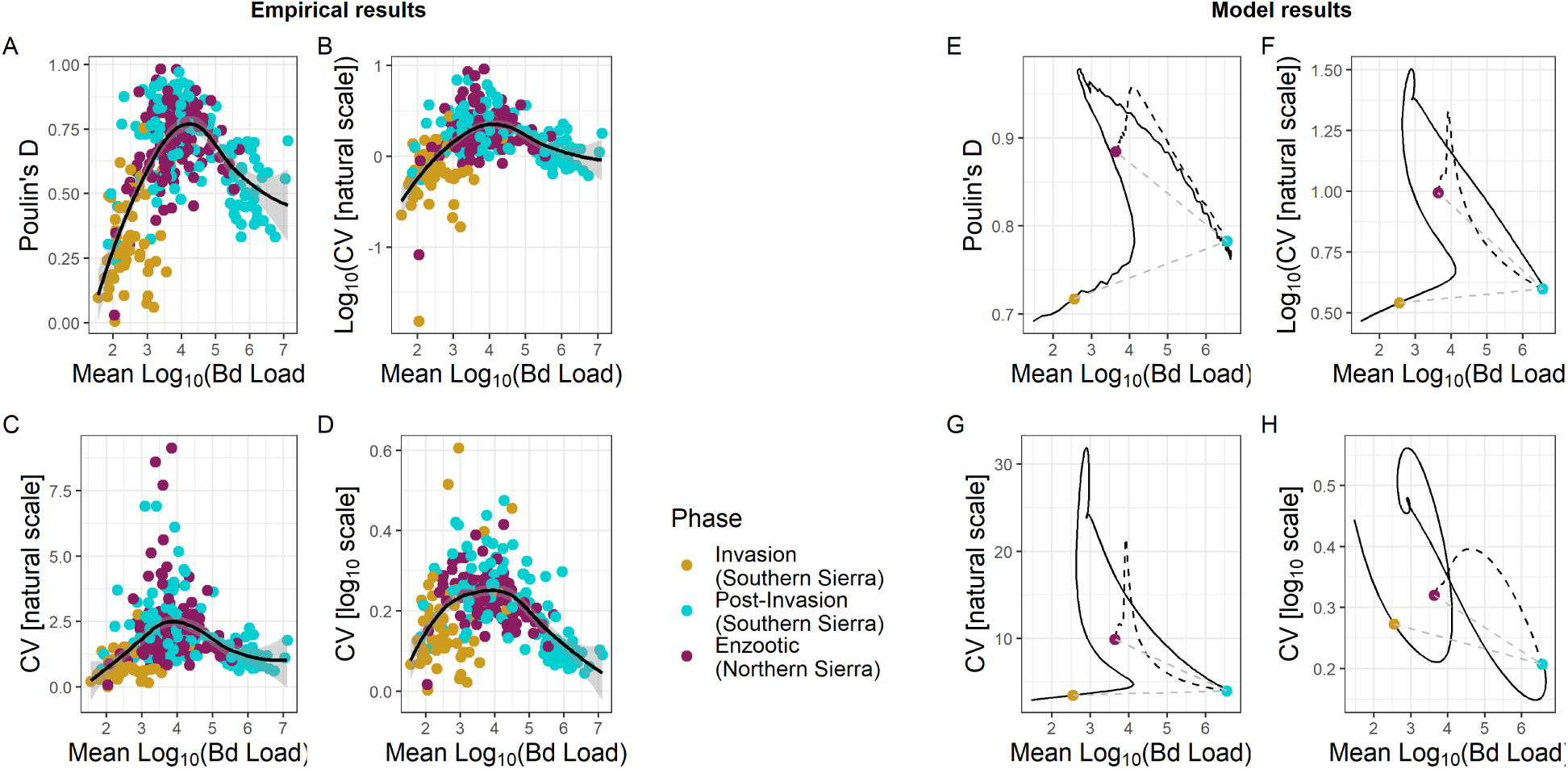
Groups in different epizoological phases plotted as a relationship between mean log_10_ Bd intensity and different aggregation metrics. **A.** Soon after Bd invasion, mean loads and aggregation (Poulin’s D) are low (yellow points). Later, post-invasion, mean loads are high and aggregation is still relatively low (blue points). Then much later, mean loads are intermediate and aggregation is higher (purple points), leading to an overall unimodal shape. This same pattern holds for other aggregation metrics including **B.** log_10_ of CV (coefficient of variation) on the natural scale, **C.** CV on the natural scale, and **D.** CV on the log_10_ scale. The unimodal trend for all empirical results is emphasized through a best-fit spline (black). The shaded gray region is the 95% confidence interval around the best fit spline. A parameterized IPM model can generally reproduce these hump-shaped patterns in all four metrics without evolution (black curve in **E-H**); we compare empirical to model results for each metric as the model need not necessarily produce a hump shape in every metric (e.g., see Fig. S5, S6). Evolution of lower pathogen growth rate (dashed black) moves populations to lower mean loads and higher aggregation metrics, generally matching the empirical results for enzootic populations. We plot the points corresponding to sampling the model results at one week (invasion, yellow), one year (post-invasion, blue), and thirty-one years (enzootic, purple) for comparison to the empirical results in Fig. 3B; low temporal resolution sampling can create a counterclockwise pattern in intensity-aggregation space (emphasized by dotted gray line connecting colored points in **E-H**).

A distinct pattern that emerges in Fig. 4A-D is the notable unimodal shape of the data in intensity-aggregation space. The downward curvature is consistent with predictions from host-macroparasite theory that intensity-dependent mortality should reduce aggregation for high mean intensities as it truncates the tail of the Bd intensity distribution resulting in lower variance for a given mean within a population. This pattern was statistically supported by strong quadratic effect of mean intensity on aggregation, with the quadratic model performing better than the linear-effect only model (ΔAIC = 150.13 from comparing a model with quadratic effect to one with only a linear relationship). Moreover, this unimodal pattern was robust to different measures of aggregation (Fig. 4A-D).

Interestingly, putative enzootic populations rarely occupy the space of high mean intensity and low aggregation (Fig. 4). We observed seven enzootic populations in this region of high mean intensity and lower aggregation. Although one group in the enzootic stage was composed of adults, the rest were subadults—a life stage that still experiences substantial disease-induced mortality even in enzootic populations [45].

#### Modeling results

Modeling showed that the unimodal intensity-aggregation patterns likely contain important, mechanistic information about disease processes. The hump-shaped patterns in intensity-aggregation space found in the field data for all four metrics did not emerge trivially from the model; depending on parameter values, the model simulations produced this hump shape for none, some, or all metrics. Simulations with parameter values based mostly on laboratory experiments [19; 47] did not produce unimodal patterns for any of the four metrics (Fig S5), indicating different biological processes may occur in the field than in a lab setting. This possibility of a quantitative mismatch between the laboratory and the field is also supported by the observation that the laboratory-based parameter values produced significantly lower values of intensity and higher values of aggregation than was observed in the field. To address this possible mismatch, we explored additional parameter sets (details in supplementary material). When we weakened the negative density dependence of pathogen growth within hosts and decreased the variance in initial infection load, our simulations produced slightly higher mean loads, lower aggregation, and a unimodal pattern in one metric, matching the field data somewhat better (Fig. S6). When we also decreased host mortality, parasite shedding rate (keeping prevalence from maxing out at one), and stochasticity in parasite growth, the model simulations produced higher intensities and still lower aggregation. Moreover, the model produced unimodal patterns for all four metrics (Fig. 4E-H). Thus, in terms of matching the intensity-aggregation patterns from the field, we considered this our best parameter set.

From host-macroparasite theory, we might expect that this unimodal pattern depends on intensity-dependent mortality driving lower aggregation at high intensity. Changing the parameter values so that hosts could survive very high loads with no mortality did increase aggregation somewhat, as expected, but unexpectedly did not significantly change the unimodal patterns (Fig. S8). Lastly, the path our simulation results take through intensity-aggregation space may explain observed counterclockwise motion through intensity-aggregation space for populations in Fig. 3B; if we sample our simulated populations at one month, one year, and thirty years to simulate the infrequent sampling of the field populations, we see how a counterclockwise motion could arise (e.g., colored points in Fig. 4E-H).

Our modeling further shows that the position of the enzootic populations in intensity-aggregation space may be a signal of host evolution. Host evolution of resistance that lowers pathogen growth rate moves populations left, toward lower mean intensity, and up, toward higher aggregation, in intensity-aggregation space for all four metrics (dashed black in Fig. 4E-H). This position of post-evolution populations higher and to the left of post-invasion populations that have experienced an epizootic but not yet evolved is consistent with the field data (Fig. 4). If hosts evolved a different defense, e.g., tolerance of higher parasite loads without dying, we would not observe this shift (Fig. S7). Thus, the enzootic populations’ position in intensity-aggregation space may indicate the evolution of resistance rather than tolerance in the host.

## Discussion

Parasite aggregation is a strong driver of disease dynamics within host populations [20]. Though aggregation in macroparasites has been extensively examined, little has been done to systematically explore aggregation within host-fungal parasite systems, despite the known impact of fungal pathogen intensity on its host. In this study, we used a dataset of nearly 57,000 samples of amphibian infection intensity to show that i) Bd is consistently more aggregated than typical macroparasites, ii) the distribution of Bd intensity within a population is generally consistent with a lognormal distribution, and iii) patterns of Bd aggregation can contain consistent signatures of biological mechanisms. This study demonstrates the utility of fungal aggregation as a means of identifying cryptic biological processes (e.g. disease-induced mortality or evolution of defense mechanisms) within host populations.

### Patterns of aggregation

Although both macroparasites and fungal parasites are aggregated within hosts, the magnitude of aggregation, as determined by the TPL slope, was significantly greater in Bd systems compared to many macroparasites. There are two possible explanations for this result that we cannot separate in this study. First, because Bd can rapidly reinfect its hosts in a process akin to within-host reproduction, we would expect levels of aggregation to be higher than most macroparasites. Supporting this expectation, [28] demonstrated similar patterns of high aggregation in *Oxyuridae* pinworms that rapidly reinfect their host (TPL slope of Oxyuridae pinworms: 2.82 [2.44, 3.22]). For both pinworms and Bd, already-infected hosts can acquire additional infection faster than uninfected hosts, increasing the variance and skew in the distribution of parasites. Second, the values of the Bd TPL slopes were highly consistent across sites and host species, indicating that levels of aggregation are conserved across populations with widely varying biology. The highly conserved nature of aggregation in macroparasites can be partially explained through statistical constraints that are independent of parasite biology [e.g., 39; 3]. This might also be true for Bd. For example, given a lognormal distribution, we would expect a TPL slope of 2, generally consistent with what we observe across Bd systems. Lognormal distributions consistently emerge in dynamic population models in ecology [32] and it is possible that the lognormal distribution of Bd (and thus the TPL slope of 2) arises because Bd dynamics, swabbing, and testing are a combination of multiplicative random processes that necessarily lead to a lognormal distribution [i.e., a central limit theorem type of argument 48]. Regardless of the exact drivers, we found that Bd aggregation does not look like that of most macroparasites.

A major impetus for quantifying patterns of aggregation in host-parasite systems is to effectively build and analyze population-level models of host-parasite dynamics. In host-macroparasite systems, the application of the negative binomial distribution has led to many basic and applied ecological insights about host-macroparasite dynamics [20; 21; 49]. Models of fungal dynamics are adopting similar approaches to those of macroparasite modeling, focusing on modeling host infection intensity [17]. However, there are still few generalizable expectations regarding fungal distributions across hosts in a population, making subsequent model assumptions somewhat tenuous.

We found that distributions of Bd intensity, conditional on infection, were most consistent with lognormal distributions across species, life stages, and locations. Lognormal distributions describe the spatial distribution of abundance and density of organisms in many natural systems and theoretically emerge in populations experiencing environmental and demographic stochasticity [32]. Surprisingly, the gamma distribution—the continuous analogue to the negative binomial distribution—performed relatively poorly when describing Bd intensity distributions. As we continue to use models to describe observed fungal infection dynamics in the field, fitting empirical data to models generally requires making some distributional assumptions about fungal intensity. Currently, host-fungal models in amphibian-Bd systems have assumed that Bd intensities are approximately lognormally distributed [50; 43], but this assumption has only been validated for a few focal amphibian-Bd systems. Our results show that a lognormal assumption is broadly applicable within amphibian-Bd systems, making theoretical and applied applications of these models robust across amphibian-Bd systems. While we only examined Bd in this study, we expect approximate lognormal distributions to hold more broadly across host-fungal systems. Testing this expectation is an important next step for uniting host-fungal dynamics under a common theoretical framework, as has been so successfully done with host-macroparasite dynamics.

### Mechanisms of aggregation

While fungal aggregation was highly consistent across amphibian species and populations, we found that there are also distinct patterns that arise in fungal aggregation that reflect underlying biological processes. In the empirical data, we observed a notable reduction in aggregation in post-invasion populations that we know were experiencing high-levels of disease-induced mortality [based on previous field observations, 15]. Moreover, we observed that the life stage in enzootic populations with the lowest levels of aggregation tended to be juveniles, the life stage in which disease-induced mortality is still occurring at a high rate even in enzootic populations [37]. While it is tempting to conclude that this pattern of reduced aggregation is solely driven by intensity-dependent mortality as predicted in host-macroparasite systems [2; 3], our modeling results show that reduced aggregation in post-invasion populations can arise even in the absence of intensity-dependent mortality.

The mechanism by which our model can produce the observed unimodal pattern in intensity-aggregation space is described as follows. When Bd first invades a population the observed intensity distribution is primarily structured by the dynamics of initial infection so that hosts have relatively similar low loads and low levels of aggregation. As the Bd outbreak proceeds, the distribution of fungal intensity begins to include both older infections with higher loads structured by within-host growth dynamics and newer infections with loads structured by initial infection dynamics. This mixture of newer and older infections increases aggregation in the intensity distribution. Most hosts become infected as the outbreak continues, and most infections are older and closer to the pathogen’s within-host carrying capacity. This drives a subsequent increase in mean intensity and reduction in aggregation. Overall, the shift from mostly newer infections to a mixture of newer and older infections then finally to mostly older infections drive a unimodal pattern in intensity-aggregation space. Our model shows that while we can get the expected unimodal pattern of reduced aggregation being driven predominantly by intensity-dependent mortality, the pattern requires that i) all hosts get infected essentially simultaneously and ii) hosts rarely lose infection during an outbreak. However, as these conditions are violated, the effect of intensity-dependent mortality on aggregation quickly becomes dwarfed by the joint effect of initial infection and within-host growth. Host-macroparasite theory has shown that there is not always a one-to-one mapping between aggregation patterns and biological processes [51]. Our results clearly highlight this point for fungal intensity distributions—there are two plausible biological mechanisms that could explain and jointly contribute to the observed reduction in aggregation at high loads: intensity-dependent mortality and the balance between initial infection and within-host growth along an epizoological trajectory. The latter is an aggregation mechanism that, to our knowledge, has not been considered in macroparasite systems, highlighting the need for unique theory describing the patterns and mechanisms of aggregation in host-fungus systems.

In addition to intensity-dependent mortality and the balance between initial infection and within-host growth, we found that patterns of aggregation contained clear signatures of the epizoological stage of a host-Bd system. Empirically, we saw populations follow a characteristic counterclockwise pattern on the yearly time scale in intensity-aggregation space. Interestingly, our modeling results illustrated that this counterclockwise pattern was likely a result of the timescale on which we observed these MYL frog-Bd systems. Our model showed that the transition from invasion phase to post-invasion epizootic phase should actually traverse a humped curve, rather than seamlessly jumping from the left to the right side of the curve. Because these sites were only sampled once a year, this data likely missed the transition from invasion phase (*<* 50% prevalence) to post-invasion phase (*>* 50% prevalence), as this often occurs rapidly within MYL frog-Bd systems. Therefore, we could only observe the invasion point and the post-invasion epizootic point within the intensity-aggregation space. Moreover, our model shows that the transition back to an intermediate intensity and high aggregation state does not occur in the model without some level of evolution in host defense; specifically, host evolution of resistance that lowers pathogen growth rate produced this pattern, while evolution of tolerance could not. MYL frog populations have persisted enzootically and begun to recover, likely due to evolved resistance to Bd [37]. As such, this is an intriguing basis for utilizing population-level aggregation patterns to identify biologically relevant processes in wild populations, such as the evolution of host defense.

Examining aggregation patterns in a system where epizoological phase was known a priori enabled us to discern patterns across the intensity-aggregation space. The full epizoological trajectory of many amphibian populations is rarely observed, and it is well known that similar amphibian populations infected with Bd can be at different places along an epizoological trajectory or on different trajectories altogether [52]. By substituting spatial replication across populations for temporal replication within populations, we show that intensity-aggregation space can help locate disparate populations along a common epizoological trajectory. We expect the approach we develop to be particularly useful for species that are generally considered to be persisting enzootically with Bd, but in reality, may be experiencing cryptic invasions and epizootics across populations (e.g., Fig. S4).

## Conclusions

Beyond amphibian-Bd systems, our study is useful for understanding fungal parasite dynamics in other wildlife populations. By extending our analyses to other host-fungal parasite systems, such as those involving white-nose syndrome in bats or *B. salamandrivorans* in amphibians, we can elucidate broader patterns of aggregation of fungal parasites. This comparative approach can unveil commonalities and distinctions in fungal intensity patterns across different hosts and parasites. Identifying patterns of aggregation and how they reflect biological processes in diverse systems has implications for conservation strategies, disease management, and disease modeling efforts. By demonstrating the ubiquity of aggregation, identifying distributional characteristics, and deciphering the biological significance of these patterns, we advance our understanding of host-fungal parasite ecology and pave the way for broader consideration of the implications of microparasite aggregation in wildlife disease ecology and epidemiological theory.

## Acknowledgements

**Personnel and other acknowledgments**

We thank T. McDevitt-Galles, W. Moss, D. Calhoun, R. Chen, T. Riepe, K. Leslie, A. Barbella, K. Rose, Dan Wetzel, Aimee Danly, Caitlin Nordheim, Miranda Kosowsky, Laura A. Brannelly, Karie A. Altman, Renato A. Martins, J. Vargas Soto, E. Hegeman, and A. Lindauer and the many other individuals who helped with data collection, processing, management, and conceptual insights.

Institutes and organizations supporting this work were: University of Pittsburgh (IACUC Protocol #1602771); Vanderbilt University (IACUC #M1600250-0); UMass (IACUC #2014003); University of California-Santa Barbara; São Paulo Research Foundation (FAPESP propc #2021/10639-5); Brazilian National Council for Scientific and Technological Development (CNPq proc #304713/2023-6); Sierra Nevada Aquatic Research Laboratory; East Bay Regional Parks and Municipal Utility Districts; Santa Clara County Parks; Sequoia-Kings Canyon and Yosemite National Parks; Inyo and Sierra National Forests; Louisiana Department of Wildlife and Fisheries (Scientific Research and Collecting Permits LNHP-17-029, LNHP-18-005, WDP-19-010); Pennsylvania Fish and Boat Commission; Tennessee Wildlife Resource Agency (Scientific Collection Permit #1546); Vermont Fish and Wildlife Department (Permit SR-2016-17); California Department of Fish and Wildlife; U.S. Fish and Wildlife Service; and multiple private landowners.

**Funding**

This project was supported by the National Park Service (to R.A.K.), Yosemite Conservancy (to R.A.K.), the US Fish and Wildlife Services Endangered Species Conservation and Recovery Grant Program, the National Science Foundation (EF-0723563, to C.J.B.; DEB-1557190, to C.J.B.; DEB-2133401, to M.Q.W.; and DBI-2120084, to C.L.R.Z., DBI-2120084, to RIBBiTR; DEB-2227340, to C.G.B.; IOS 2303908, to C.G.B.; DEB-2133401, to M.Q.W; DEB-2133399, to T.C.S; DEB-1149308; DEB-1754171), and the NIH/NSF Ecology and Evolution of Infectious Diseases program (R01GM109499 and R01GM135935).

