## Supplemental Material for "Do fungi look like macroparasites? Quantifying the patterns and mechanisms of aggregation for host-fungal parasite relationships"

### Supplementary Material

#### I. Supporting Figures & Tables

Table S1: The four datasets were divided into groups based on site, sampling year, season, species, and life stage. Invasion phase was only available for the Sierra dataset. Multi-seasonal data collection primarily only took place within the Eastern US and Brazil datasets. The host species that made up at least 70% of the groups for each dataset are listed. Characteristics of the datasets are a result of sampling effort and not true demography of the host populations.

| **Datasets** | | | | |
|---|---|---|---|---|
| | **Brazil** | **East Bay** | **Eastern US** | **Sierra** |
| Years collected | 2018-2023 | 2013-2017 | 2016-2019 | 2004-2019 |
| Sampling seasons | Wet(Nov-Apr), Dry(May-Oct) | Summer (May-Sep) | Winter, Spring, Summer, Fall | Summer (Jun-Sep) |
| Sampled life stages | Larva, Adult | Larva, Subadult, Adult | Larva, Subadult, Adult | Larva, Subadult, Adult |
| Number of sites | 35 | 169 | 31 | 620 |
| Number of records | 4,365 | 10,490 | 12,457 | 29,600 |
| **Number of Groups** | | | | |
| Poulin's D & CV analysis (min.2 Bd-positive) | 59 | 326 | 257 | 444 |
| TPL analysis (min.3 Bd-positive) | 49 | 267 | 231 | 414 |
| Distribution fit (min. 10 Bd-positive) | 5 | 81 | 110 | 329 |
| *continued on next page* | | | | |

| TPL Analysis Groups (min. 3 Bd-positive) | | | | | | | | |
|---|---|---|---|---|---|---|---|---|
| **Life Stage** | | | | | | | | |
| Larva | 1 | 2.0% | 47 | 17.6% | 1 | 0.4% | 123 | 29.7% |
| Subadult | 0 | 0.0% | 192 | 71.9% | 49 | 21.2% | 85 | 20.5% |
| Adult | 5 | 10.2% | 14 | 5.2% | 166 | 71.9% | 206 | 49.8% |
| Unknown | 43 | 87.8% | 14 | 5.2% | 15 | 6.5% | 0 | 0.0% |
| **Invasion Phase** | | | | | | | | |
| Epizootic - Invasion | *NA* | | *NA* | | *NA* | | 41 | 9.9% |
| Epizootic - Post Invasion | *NA* | | *NA* | | *NA* | | 212 | 51.2% |
| Enzootic | *NA* | | *NA* | | *NA* | | 161 | 38.9% |
| **Season** | | | | | | | | |
| Winter | 0 | 0.0% | 0 | 0.0% | 19 | 8.2% | 0 | 0.0% |
| Spring | 0 | 0.0% | 0 | 0.0% | 110 | 47.6% | 0 | 0.0% |
| Summer | 0 | 0.0% | 267 | 100.0% | 73 | 31.6% | 414 | 100.0% |
| Fall | 0 | 0.0% | 0 | 0.0% | 29 | 12.6% | 0 | 0.0% |
| Dry | 24 | 49.0% | 0 | 0.0% | 0 | 0.0% | 0 | 0.0% |
| Wet | 25 | 51.0% | 0 | 0.0% | 0 | 0.0% | 0 | 0.0% |
| **Host Species** | | | | | | | | |
| *Anaxyrus boreas* | 0 | 0.0% | 53 | 19.9% | 0 | 0.0% | 0 | 0.0% |
| *Brachycephalus pitanga* | 22 | 44.9% | 0 | 0.0% | 0 | 0.0% | 0 | 0.0% |
| *Hylodes phyllodes* | 4 | 8.2% | 0 | 0.0% | 0 | 0.0% | 0 | 0.0% |
| *Ischnocnema henselii* | 9 | 18.4% | 0 | 0.0% | 0 | 0.0% | 0 | 0.0% |
| *Pseudacris crucifer* | 0 | 0.0% | 0 | 0.0% | 33 | 14.3% | 0 | 0.0% |
| *Pseudacris regilla* | 0 | 0.0% | 152 | 56.9% | 0 | 0.0% | 0 | 0.0% |
| *Rana catesbeianus* | 0 | 0.0% | 2 | 0.7% | 40 | 17.3% | 0 | 0.0% |
| *Rana clamitans* | 0 | 0.0% | 0 | 0.0% | 62 | 26.8% | 0 | 0.0% |
| *Rana muscosa/sierrae* | 0 | 0.0% | 0 | 0.0% | 0 | 0.0% | 444 | 100.0% |
| *Rana sphenocephalus* | 0 | 0.0% | 0 | 0.0% | 27 | 11.7% | 0 | 0.0% |
| *Taricha torosa* | 0 | 0.0% | 32 | 12.0% | 0 | 0.0% | 0 | 0.0% |

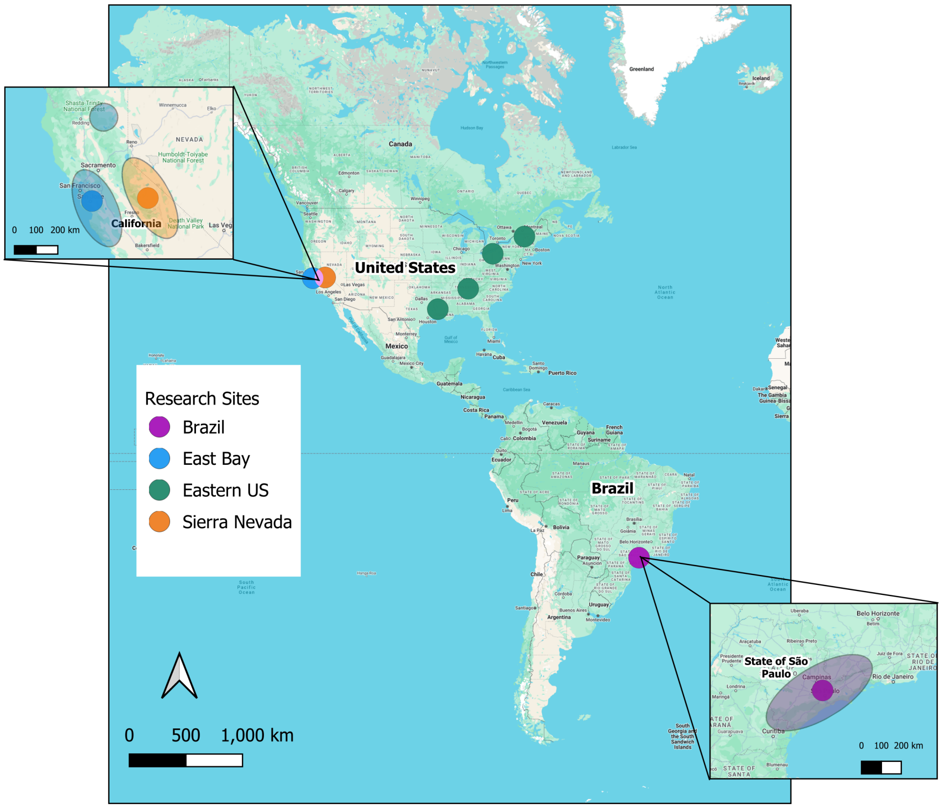

Figure S1: Map of the sampling sites for the four datasets: Eastern US (green), Sierra (orange), East Bay (blue), and Brazil (purple). Insets magnify the study regions in California and Brazil. The shaded ellipses provide the general extent of the sampling locations, color-coded by dataset.

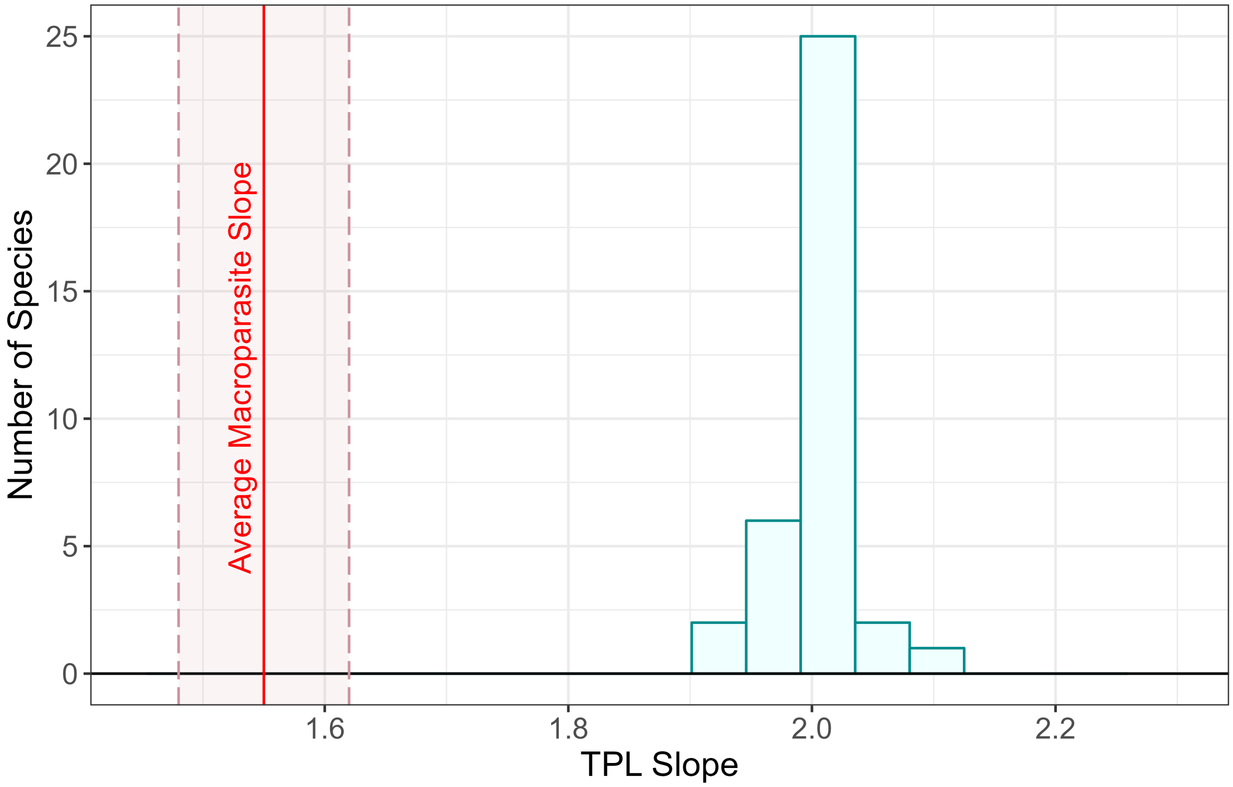

Figure S2: The distribution of species-specific Taylor's Power Law slopes. All are distinctly greater than the slope which is typically seen in macroparasite systems (mean slope [red] and 95% CI [shaded pink]).

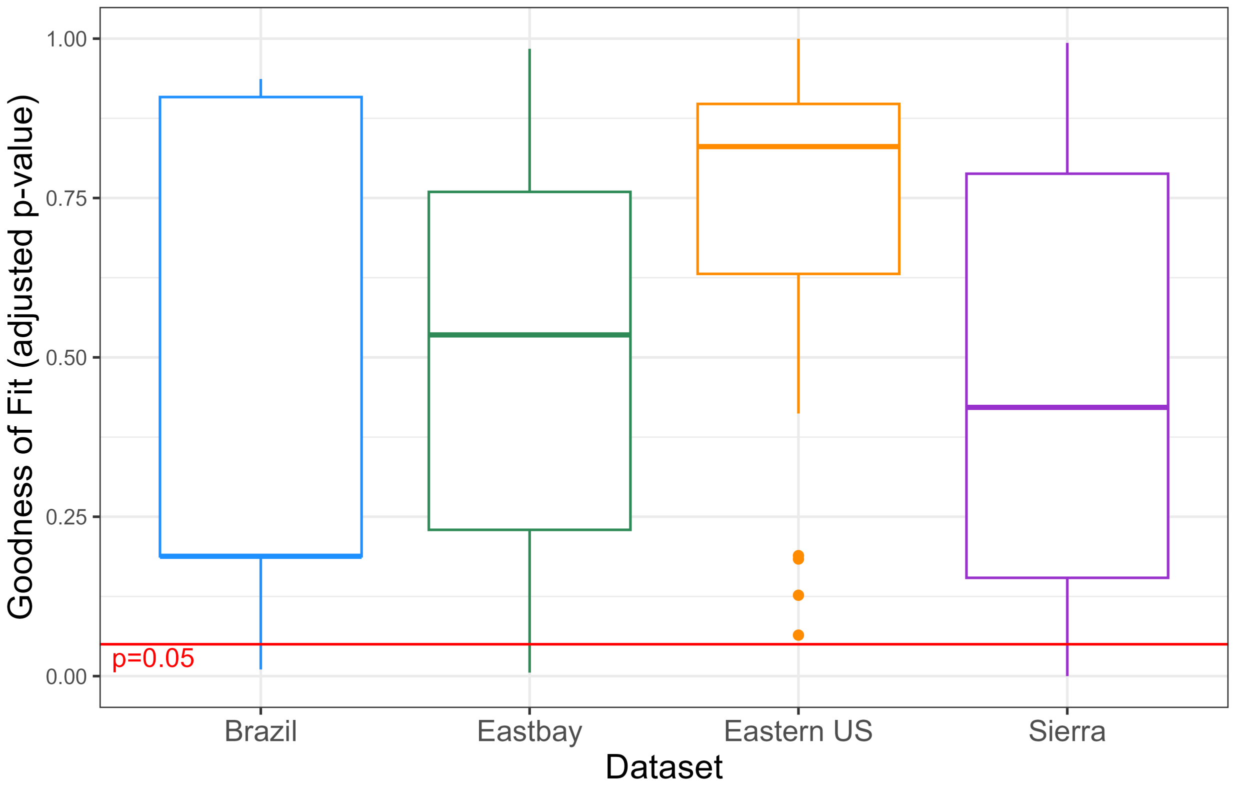

Figure S3: Adjusted p-value results of the Shapiro-Wilks tests on the log-transformed fungal loads for all the groups across the data sets.

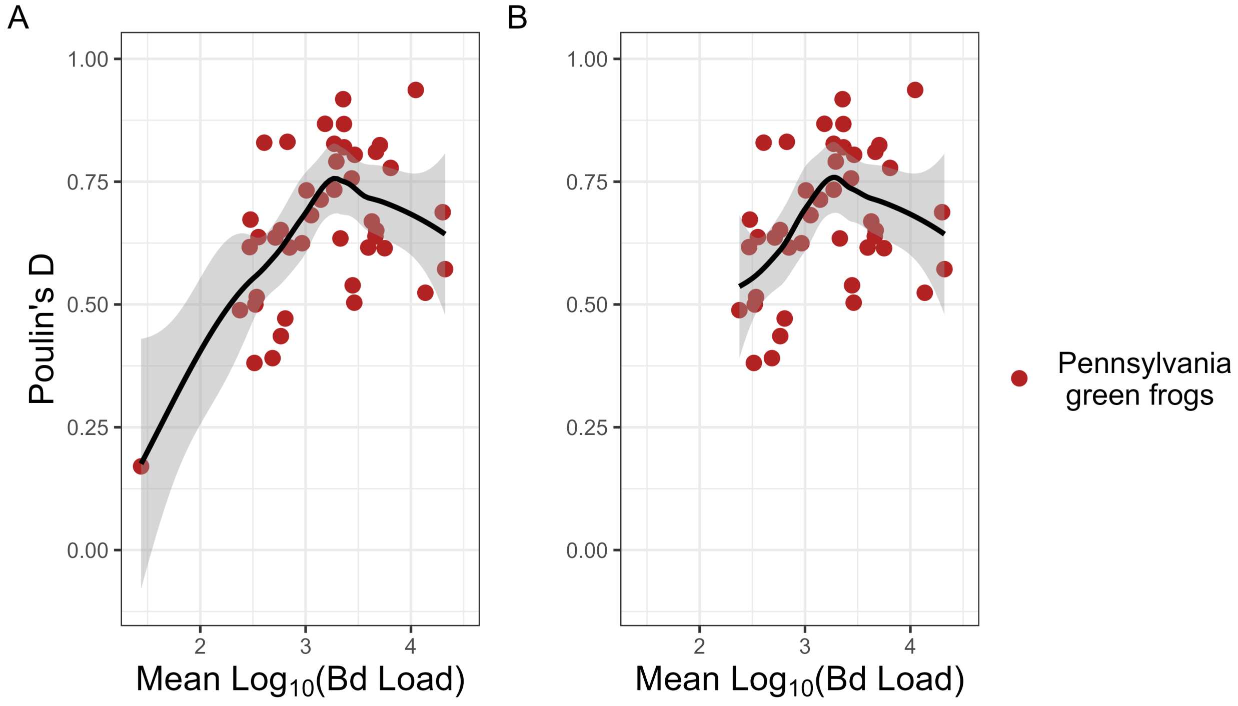

Figure S4: **A.** Groups of green frogs (*Rana clamitans*) in Pennsylvania plotted in intensity-aggregation space. *Rana clamitans* (green frogs) are generally considered to robustly persist in an enzootic state with Bd in the eastern US [e.g., 1]. A trend line (black) and corresponding 95%CI (gray) demonstrate a pattern consistent with what may be expected when host populations are experiencing a range of epizoological phases—invasion [perhaps better categorized as a "spillover host phase" in this case, 2], post-invasion epizootic, and enzootic. **B.** This general pattern remains even when the one group most consistent with the invasion phase is removed, demonstrating that this single low-load, low-aggregation group is not solely driving the overall trend.

### II. Integral Projection Model

We developed an Integral Projection Model (IPM) that tracks the distribution of log Bd load. The IPM is similar to a previously published IPM [3] but we add host birth and evolution. The model tracks the distribution of ln(pathogen load) = $x$ at discrete time point, $t$. In a one-day time step, susceptible hosts survive with some probability $s_0$ and become infected with a probability that depends on the transmission rate parameter, $\beta'$, and the number of pathogens in the environment, $Z$. The probability that a new infection starts with load $x$ follows normal probability density $G_0(x)$ with some mean ($\mu_0$) and standard deviation ($\sigma_0$). Infected hosts have some probability of recovery in a time step that decreases as $x$ increases [$l(x) = 1-(1-e^{y+wx}/(1+e^{y+wx}))^{1/3}$ with $w < 0$ (see Table S2 for parameter definitions); this formulation reflects changing from the three-day time step this recovery process was empirically parameterized with to our one-day time step]. Infected hosts also have a probability of survival for a time step, $s(x)$, that declines with $x$ [precisely, $s(x) = 1 - \phi((x - \mu_{LD50})/\sigma_{LD50})$ where $\phi$ is the cumulative distribution function of the standard normal and $\sigma_{LD50} > 0$]. Infected hosts that survive and do not recover can experience a decrease or increase in load so that the probability of a given host having load $x'$ in the next time step follows a normal density as a function of the current $x$ (precisely, $G(x'|x)$ is a normal density with mean $a + bx$ and fixed standard deviation $\sigma_G$). The form of the growth function is such that mean log load increases linearly at low $x$ then more slowly as $x$ increases to a within-host carrying capacity of $a/(1-b)$, neglecting death and recovery (assuming $a > 0$, $1 > b > 0$). Infected hosts produce $\lambda$ environmental pathogens for every pathogen they have (total shed is $\lambda e^x$, because the pathogen in the environment is modeled on the linear scale) and environmental pathogens have a probability $\nu$ of persisting in the environment. We add host reproduction and multiple genotypes so we can simulate evolution.

We consider how host genotypes may differ, leading to host evolution. We do not have strong *a priori* expectations for inheritance, etc. Therefore, we model host evolution simply as competition among a fixed number ($n = 10$ in our simulations) of asexually reproducing genotypes where a genotype $j$ may differ from other genotypes in fecundity or defense against pathogens (parameters with a $j$ subscript are considered as differing by genotype). The total number of hosts of a genotype ($N_j$, summing across infection status and loads) reproduce with some maximum, per-capita fecundity $r_j$, $m$ giving the probability of mutation so that reproduction produces a genotype other than the parental genotype, and crowding that reduces fecundity in a density-dependent manner; crowding is parameterized by $q$ and we constrain fecundity to be non-negative (Eq. 1a but we do not consider high enough densities to drive negative fecundity even if we did not constrain fecundity; note that births of genotype $j$ is a function of the density of all genotypes due to mutation and crowding). Thus, an isolated population of host genotype $j$ without disease grows logistically toward a carrying capacity $[(r_j + s_0 - 1)/(r_j q)]$. Tolerance can vary by genotype by varying $\mu_{LD50,j}$ and resistance to pathogen growth can vary by genotype by varying $a_j$. We leverage the large amount of data on our focal frog system to parametrize this model with parameter values that are reasonable for the laboratory [3, see Table S2 for full list of values and supplementary figure captions for alternate parameter values we explored].

We used this model to simulate epizootics and eco-evolutionary processes. Initially, simulations began with wild-type hosts at their disease-free equilibrium and a single environmental parasite per cubic meter. We ran these epizootic simulations for one year to consider what patterns would arise without host evolution. Then we considered the effect of host evolution by simulating host evolution after that single year of epizootic; the wildtype genotype began eco-evolutionary simulations at the same density and distribution of load for which it ended the epizootic simulation, as did environmental parasites. Genotypes other than the wildtype began with the same infection prevalence and distribution of load as the wildtype but density lower than the wildtype's by a factor of $10^{-6}$. These other genotypes (we chose $n = 10$ genotypes total) differ from the wild-type in that they have a fitness advantage in terms of some trait that provides defense against pathogens ($y_j$ ranging from the least defended, wild-type genotype, $y_{wt}$, to the most defended genotype, $y_{defended}$); genotypes pay a fecundity cost of this defense $r_j = r_{wt} - (r_{wt} - r_{defended})[(y_{wt} - y_j)/(y_{wt} - y_{defended})]^w$. Thus, the wild-type genotype has the highest fecundity ($r_{wt}$); the fecundity cost per unit of defensive trait is small, at first, then accelerates rapidly driving a sudden decline in fecundity to the lowest fecundity of the most defended genotype ($r_{defended}$). These accelerating costs (accelerating because we chose $w > 1$, specifically, $w = 10$), generally select for an intermediate degree of host defense [4]. We ran eco-evolutionary simulations for thirty years. All model simulations were performed in R.

Eq. S1a: Birthrate of genotype $j$

$$f_j(N_{1:n}(t)) = [\underbrace{r_j N_j(t)(1-m)}_{\text{Birth less mutation}} + \underbrace{\frac{m}{n-1}\sum_{k=1,k\neq j}^{n} r_k N_k(t)\,]}_{\text{Gains from mutation}} \underbrace{max(0, 1 - qN_t(t)))}_{\text{Crowding}}$$

Eq. S1b: Susceptible individuals of genotype $j$

$$S_j(t+1) = \underbrace{f_j(N_{1:n}(t))}_{\text{Birth}} + \underbrace{S_j s_0 e^{-\beta' Z(t)}}_{\text{S}_\text{j} \text{ survive, remain uninfected}} + \underbrace{\int_{-\infty}^{\infty} I_j(x,t) s_j(x) l(x) dx}_{\text{I}_\text{j} \text{ survive, recover}}$$

Eq. S1c: Infected individuals of genotype $j$

$$I_j(x', t+1) = \underbrace{S_j s_0 [1 - e^{-\beta' Z(t)}] G_0(x')}_{\text{S}_\text{j} \text{ survive, get infected}} + \underbrace{\int_{-\infty}^{\infty} I_j(x,t) s_j(x)(1 - l(x)) G_j(x'|x) dx}_{\text{I}_\text{j} \text{ survive, do not recover}}$$

Eq. S1d: Environmental Bd

$$Z(t+1) = \underbrace{\lambda \int_{-\infty}^{\infty} e^x \sum_{j=1}^{n} I_j(x,t) dx}_{\text{Bd shedding}} + \underbrace{\nu Z(t)}_{\text{Environmental persistence}}$$

Table S2: Parameter values, units, and sources. Parameters with a "wt" in their subscript are allowed to vary by host genotype for some evolution scenarios; the wildtype value is listed. SD abbreviates standard deviation.

| **Parameter** | **Value and units** | **Source** |
|---|---|---|
| $r_{wt}$: Offspring per host per day | 0.171 | Assuming an egg mass of 200 eggs per year [5] but dividing across both sexes and a representative duration of the lifespan spent as an adult [7.5 yrs; [6; 7]] compared to as a metamorph [2.5 yrs; [7]] and tadpole [2 yrs; [6]] |
| $q$: Negative density-dependence of host fecundity | 0.998 $m^3$ $host^{-1}$ | Chosen to give an equilibrium host density of 1 hosts $m^3$ without disease |
| $m$: Probability of mutation to a different host genotype | $5 \times 10^{-6}$ | We take a high mutation rate measured in another amphibian species [8] as an upper limit and assume a $1 \times 10^{-3}$ probability that a mutation changes the host to another genotype. |
| $1 - s_0$: Probability of an uninfected host's death per day | $2.282 \times 10^{-4}$ | Corresponds to an average lifespan of 12 years in the absence of disease [6; 7]. |
| $\beta'$: Transmission rate | $4.035 \times 10^{-5}$ | Corresponds to the value fitted to transmission rate data by [9]. |
| $\mu_0$: Mean of initial log load | 3.382 log pathogens | Found by [3] and choosing $20°C$ as a representative temperature. |
| $\sigma_0$: SD of initial log load | 2.708 log pathogens | Found by [3] and choosing $20°C$ as a representative temperature. |
| $\alpha_{wt}$: Log pathogen growth rate in one day | 0.664 log pathogens | Found by [3] and choosing $20°C$ as a representative temperature and converting from the three-day time step used to our one-day time step. |
| $b$: Density-dependence of within-host pathogen growth | 0.928 | Found by [3] and choosing $20°C$ as a representative temperature and converting from the three-day time step used to our one-day time step. |
| $\sigma_G$: SD of pathogen growth function | 1.124 log pathogens | Found by [3] and choosing $20°C$ as a representative temperature, assuming 6 as a representative log load and converting from the three-day time step used to our one-day time step. |

| | | |
|---|---|---|
| $\mu_{LD50,wt}$: $LD_{50}$ of the host survival function | 15.949 log pathogens | From matching survival curve used by [3] and converting from the three-day time step used to our one-day time step. |
| $\sigma_{LD50,wt}$: Shape parameter of host survival function | 2.953 log pathogens | From matching survival curve used by [3] and converting from the three-day time step used to our one-day time step. |
| $y$: Governs how load affects the probability of recovery at load $x = 0$ | -1.807 corresponds to 0.049 probability | Found by [3] and choosing $20°C$ as a representative temperature. |
| $w$: Governs how load affects the probability of recovery | -0.472 corresponds to 0.049 probability | Found by [3] and choosing $20°C$ as a representative temperature. |
| $\lambda$: Bd shedding rate per day | 0.06 | Not well known but adjusted so that parasites spread during the epizootic but not to 100% prevalence. |
| $\nu$: Pathogen survival probability in the environment per day | $9.482 \times 10^{-3}$ | From the total encystment and mortality rate of zoospores found by [10] at $23°C$. |

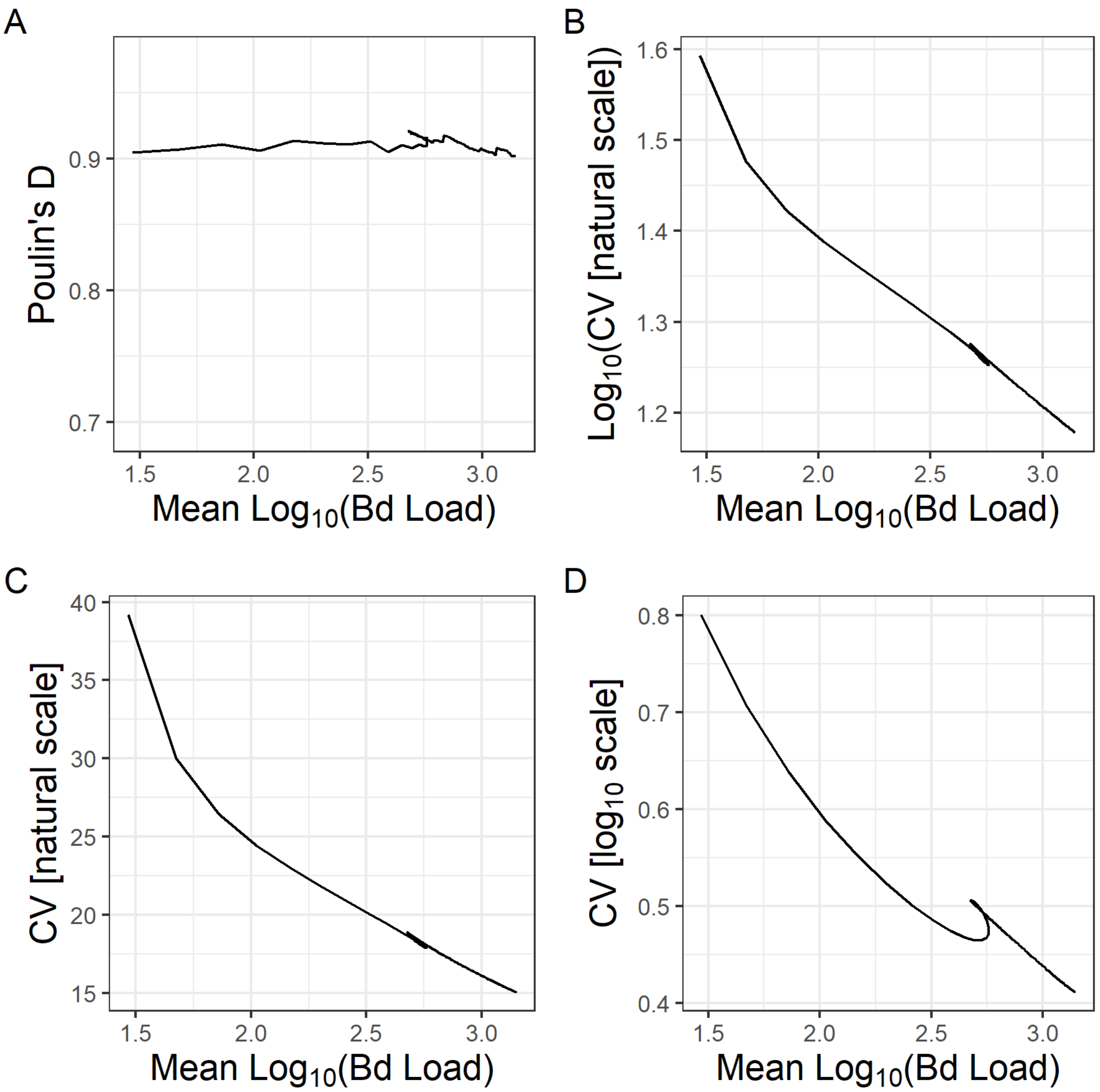

Figure S5: Simulations from laboratory-based parameter values. We calculate mean $\log_{10}$ Bd load and all four aggregation metrics across ($\log_{10}$ where log is used) from Bd invasion for one year. Hump-shaped patterns do not emerge in any metric.

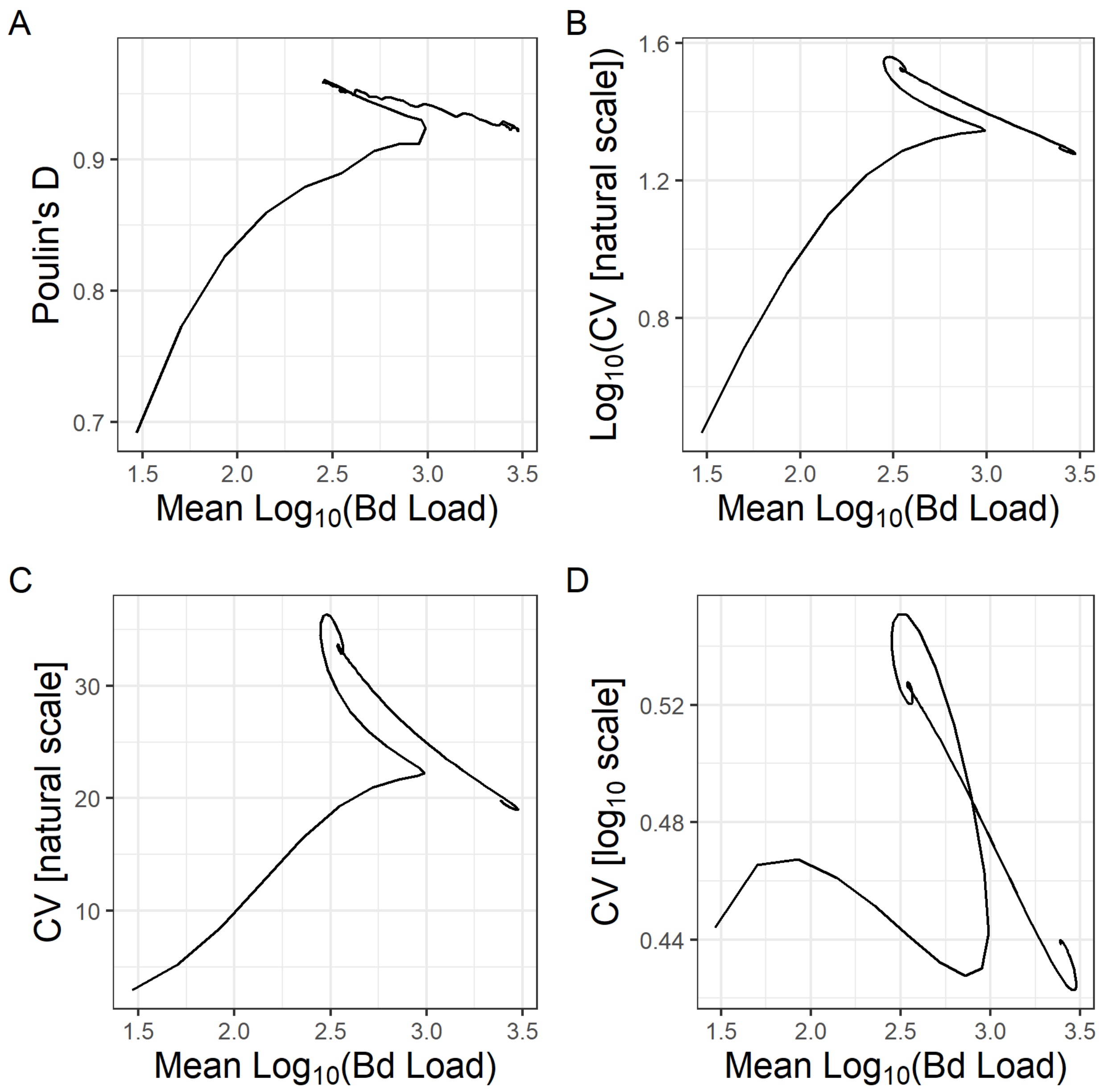

Figure S6: Simulations from slightly altered parameter values. Compared to the laboratory parameter values, we decreased negative density dependence of within-host growth (raised $b$ from 0.928 to 0.960) and lowered the variation in infection load upon infection (lowered $\sigma_0$ from 2.708 to 1.50). We found these parameter changes improved match to the field data through increased mean loads, decreased aggregation metrics (**A-D**; compared to Fig. S5), and a hump-shaped pattern in CV on the $\log_{10}$ scale (D).

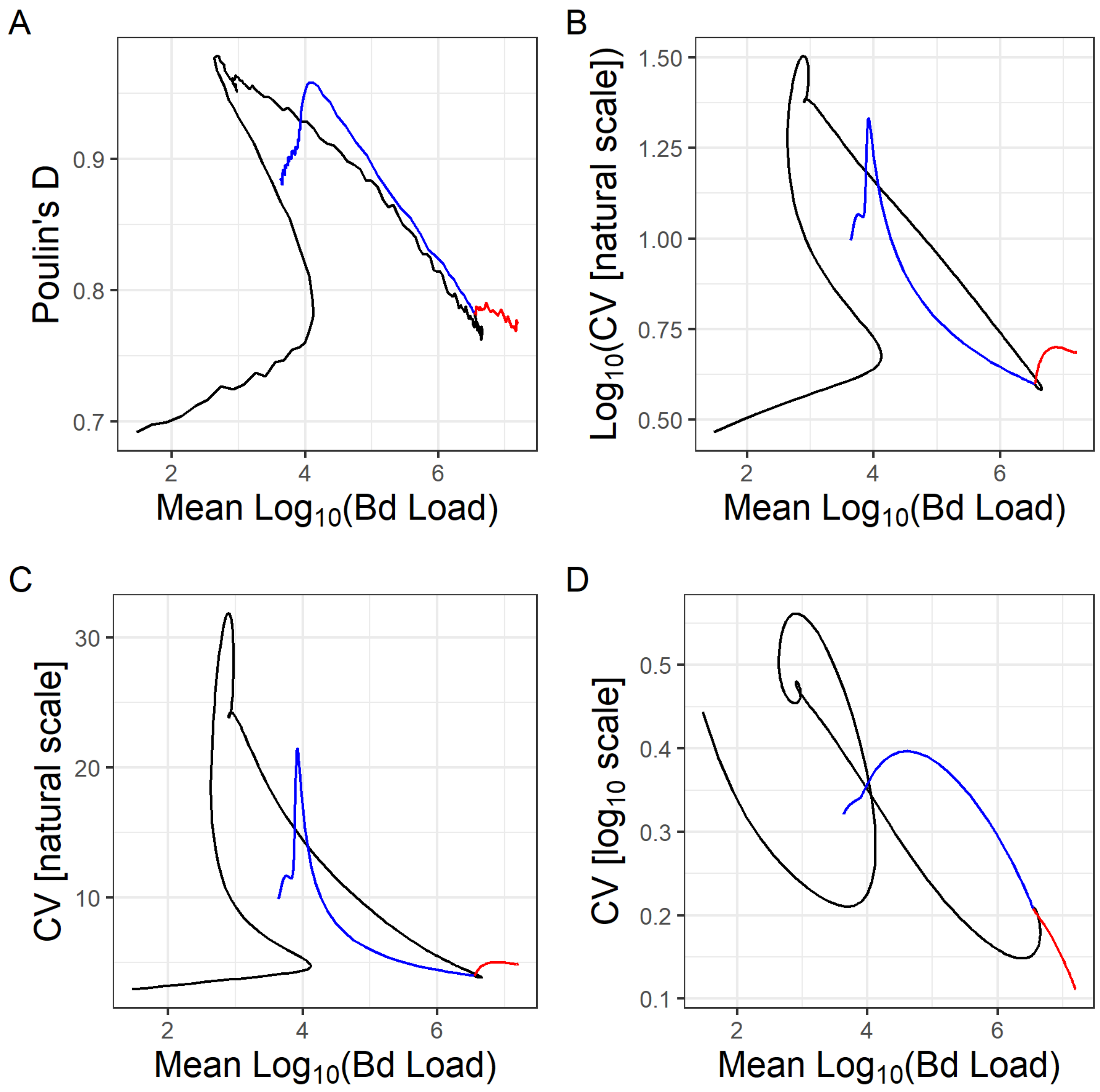

Figure S7: Compared to the laboratory parameter values, we decreased negative density dependence of within-host growth (raised $b$ from 0.928 to 0.960), lowered the variation in infection load upon infection (lowered $\sigma_0$ from 2.708 to 1.50), decreased the death rate of infected hosts (raised $\mu_{LD50}$ from 15.949 to 25), decreased stochasticity in parasite growth (lowered $\sigma_G$ from 1.124 to 0.5), and decreased parasite shedding from infected hosts (lowered $\lambda$ from 0.06 to 0.015). We found these parameter changes improved match to the field data through increased mean loads, decreased aggregation metrics (**A-D**; compared to Fig. S5), and a hump-shaped pattern in all aggregation metrics. After one year of simulation without host evolution (black curve), we simulated 30 years with evolution through competition of 10 host genotypes that either varied in resistance to parasite growth ($a$, blue curve) or tolerance of parasite loads ($\mu_{LD50}$, red curve).

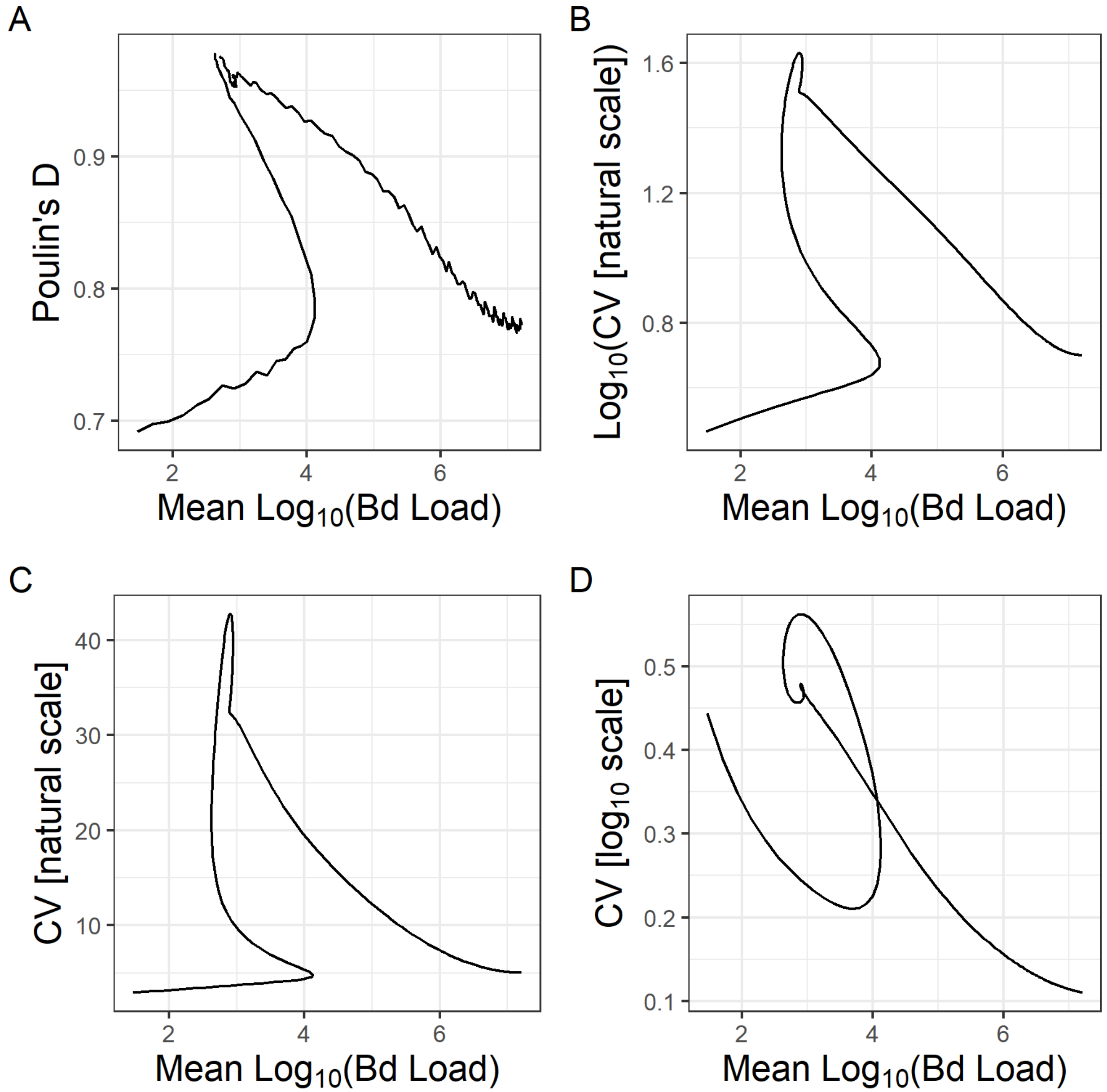

Figure S8: Intensity-dependent host death is not required for hump-shaped patterns in intensity-aggregation space. We kept all parameters the same as in Fig. S7 except we essentially eliminated all death of infected hosts (raised $\mu_{LD50}$ from 25 in Fig. S7 to $10^4$). This increased aggregation somewhat, most notably $\log_{10}$(CV natural scale) and CV on the natural scale (compared to Fig. S7) but did not substantially alter any of the hump-shaped patterns.
